# Co-Increasing Neuronal Noise and Beta Power in the Developing Brain

**DOI:** 10.1101/839258

**Authors:** Wei He, Thomas Donoghue, Paul F Sowman, Robert A Seymour, Jon Brock, Stephen Crain, Bradley Voytek, Arjan Hillebrand

## Abstract

Accumulating evidence across species indicates that brain oscillations are superimposed upon an aperiodic 1/*f* - like power spectrum. Maturational changes in neuronal oscillations have not been assessed in tandem with this underlying aperiodic spectrum. The current study uncovers co-maturation of the aperiodic component alongside the periodic components (oscillations) in spontaneous magnetoencephalography (MEG) data. Beamformer-reconstructed MEG time-series allowed a direct comparison of power in the source domain between 24 children (8.0 ± 2.5 years, 17 males) and 24 adults (40.6 ± 17.4 years, 16 males). Our results suggest that the redistribution of oscillatory power from lower to higher frequencies that is observed in childhood does not hold once the age-related changes in the aperiodic signal are controlled for. When estimating both the periodic and aperiodic components, we found that power increases with age in the beta band only, and that the 1/*f* signal is flattened in adults compared to children. These results suggest a pattern of co-maturing beta oscillatory power with the aperiodic 1/*f* signal in typical childhood development.

## INTRODUCTION

### Neuronal Oscillations Characteristic of Childhood Brain Maturation

Neuronal oscillatory power undergoes profound developmental changes throughout childhood (Gomez et al., 2017; Rodriguez-Martinez et al., 2017). These developmental changes in neuronal oscillations are often assessed noninvasively using electrophysiological brain recordings such as magneto-/electro-encephalography (MEG/EEG). Commonly, a Fourier analysis is used to compute the power spectral density (PSD) in fixed frequency bands, including delta (1-4 Hz), theta (4-8 Hz), alpha (8-13 Hz), beta (13-30 Hz), and gamma (30-80 Hz; Mackay, 1997; note that different studies may differ slightly in establishing the boundaries between frequency bands). There are age-related decreases in total power, which is estimated across a broad frequency range (Gasser et al., 1988; Schafer et al., 2014; Gomez et al., 2017; Rodriguez-Martinez et al., 2017), as well as age-related decreases in absolute power in each of the narrowband frequencies (Gasser et al., 1988; Boord et al., 2007). Studies also report a low-to-high redistribution of relative power (i.e., where power is estimated in any given band in relation to the total power across all frequencies); more specifically, relative power decreases in the delta and theta bands and increases in the alpha, beta and gamma bands (Puligheddu et al., 2005; Gomez et al., 2013; Schafer et al., 2014). In addition, there is an increase during childhood in the peak frequency of alpha oscillations, which typically reach a peak frequency of ∼10 Hz around the primary/elementary school years (Marcuse et al., 2008; Boersma et al., 2011; Cragg et al., 2011; Smit, Boomsma, et al., 2012; Miskovic et al., 2015; Gomez et al., 2017; Rodriguez-Martinez et al., 2017).

Power in each narrow frequency band has been associated with different cognitive functions. The alpha rhythm (Markand, 1990) has been prominently associated with inhibition of visual attention (Jensen and Mazaheri, 2010; Clayton et al., 2017; Voytek et al., 2017). In addition, the increase of alpha peak frequency with age has been considered a biomarker for cognitive development (Marcuse et al., 2008; Boersma et al., 2011; Cragg et al., 2011; Smit, Boomsma, et al., 2012; Miskovic et al., 2015; Gomez et al., 2017; Rodriguez-Martinez et al., 2017), which suggests that the perception of visual stimuli would also improve with age (Thut et al., 2012). In the sensory motor cortex, mu, which is analogous to the alpha band (Mackay, 1997), and beta oscillations have been found to increase when cortical motor areas are disengaged (Ritter et al., 2009; Jenkinson and Brown, 2011). Our recent longitudinal MEG study of motor development in children demonstrated linear increases in amplitude and mean frequency in movement-evoked mu and beta oscillations (Johnson et al., 2019). In studies of developmental resting-state neuronal activity (i.e., in the absence of any specific cognitive event), there are also reports of age-related increases in mu power from infancy to age 5 (Berchicci et al., 2011) and age-related increases in beta power between age 9-14 and 20-42 (Heinrichs-Graham et al., 2018). Delta power has been correlated with different stages of sleep (Amzica and Steriade, 1998), and theta activity has been shown to relate to executive attention and working memory (Wang et al., 2005). Developmental trends in delta and theta power are less clear, with some studies reporting profound decreases during childhood (Schafer et al., 2014; Gomez et al., 2017) and others reporting no changes between 9 and 11 years, followed by decreases into early adulthood (Campbell and Feinberg, 2009). There is evidence of increased gamma activity in the initiation and cessation of movement (Gaetz et al., 2010; Burianova et al., 2013; Cheyne and Ferrari, 2013; Marstaller et al., 2014; Sowman et al., 2014). Gamma activity can be adult-like as early as 3 years of age in motor tasks (Johnson et al., 2019), and resting state gamma activity across the first 3 years of life is predictive of later development of language and cognitive skills (Benasich et al., 2008). Based on the previous literature, it is clear that a precise characterisation of developmental changes in neuronal oscillatory power is critical for our mechanistic understanding of the maturation of cognitive functions during childhood.

### Outstanding Questions

Spectral analysis of resting-state MEG/EEG recordings has proven to be a powerful tool for assessing age-related power changes. However, nearly all previous resting-state studies have used the five canonical frequency bands to estimate developmental changes in power. These studies, therefore, are susceptible to methodological challenges. For example, examining power in pre-defined bands can conflate power changes with other parameters, such as oscillation centre frequency and bandwidth (Haller et al., 2018). An example is the increase in alpha peak frequency with age, which is considered to be one of the most important electrophysiological hallmarks of brain development (Valdés et al., 1990). Following historical tradition, nearly every study to date has defined the peak frequency as the frequency with the highest amplitude within the range of the canonical alpha band. Under this constraint, it is impossible to determine whether or not changes in the peak frequency may in fact reflect shifts of the peak frequency outside the canonical alpha band. Similarly, group-level estimates of power in one band may leak into the estimates of power in adjacent bands, given the variability in oscillation centre frequency across individuals (Haegens et al., 2014; Samaha and Postle, 2015) and age (Rodriguez-Martinez et al., 2017). But perhaps most importantly, the narrowband estimates of oscillatory activity may be affected by aperiodic components in the signal. Brain activity at many spatiotemporal scales, ranging from neuronal membrane potentials to MEG/EEG signals, exhibits an aperiodic background signal that co-exists with neuronal oscillations (He, 2014). This aperiodic signal follows a power-law function: *P* ∝ 1/*f*^x^. Thus, power *P* is inversely proportional to frequency *f* with a power-law exponent of (, which is equivalent to the slope of the power spectrum when plotted in the log-log space. The aperiodic 1/*f* signal is not only prevalent in the nervous system, but it is a ubiquitous feature of a wide variety of time-varying real-world systems, including the flow of the river Nile and the luminosity of stars (Bak et al., 1987). Historically, the aperiodic 1/*f* signal has not been as well investigated as periodic (oscillatory) brain activity, resulting in a lack of consensus on how to measure it, what it may reflect, and what might be its physiological generators. Nonetheless, the recent electrophysiological literature has started to elucidate the functional significance of the 1/*f* signal in human cognition and behaviour. For instance, this signal has been found to vary systematically with age (Voytek et al., 2015), and to change with task demands (He et al., 2010) that can co-vary with behavioural performance (Podvalny et al., 2015). Moreover, recent simulations, as well as empirical data from rats and macaques, indicate that neuronal excitation and inhibition in cortical circuits can be inferred from the slope of the invasively-recorded electrophysiological power spectrum (Gao et al., 2017). This further emphasises the importance of quantifying the 1/*f* signal and its contributions to human cognition (Voytek and Knight, 2015).

Related to this, standard analytic approaches, which estimate power in narrow frequency bands, fail to examine whether an oscillation – a rhythmic component that peaks at a particular frequency – is truly present in the power spectrum. The power that is assessed within a pre-defined frequency range of the electrophysiological power spectrum is most likely a mixture of both oscillatory and aperiodic 1/*f* components. It is imperative to disentangle age-related changes in the narrowband oscillations from those in the broadband aperiodic 1/*f* signal. Although these signals are inter-related, they are likely to represent distinct underlying neural mechanisms (Haller et al., 2018).

### Current Study: Aims and Hypotheses

To overcome the limitations of previous studies that used narrowband power analyses, the present study used advanced analysis techniques (Haller et al., 2018) to investigate, and to disentangle, age-related effects in the aperiodic 1/*f* and the oscillatory components of brain activity. For this, we used a paediatric whole-head MEG scanner (Johnson et al., 2010) to collect resting-state electrophysiological signals from children ranging in age from 4 to 12 years, as well as a conventional MEG scanner to obtain the same signals from adults. Importantly, source waveforms were computed using an atlas-based beamforming approach for power spectra analyses (Hillebrand et al., 2012; Hillebrand et al., 2016), which allowed for direct comparison of MEG data acquired from the two systems (He et al., 2019).

We first analysed neuronal power in pre-defined frequency bands using standard methods, to permit direct comparisons of our results with those from previous studies. In line with the previous literature (Gasser et al., 1988; Puligheddu et al., 2005; Marcuse et al., 2008; Boersma et al., 2011; Cragg et al., 2011; Smit, Boersma, et al., 2012; Gomez et al., 2013; Schafer et al., 2014; Miskovic et al., 2015; Gomez et al., 2017; Rodriguez-Martinez et al., 2017), we hypothesised that, in contrast to children, adults would show decreased low-frequency power, increased high-frequency power, and an increased alpha peak frequency.

Subsequently, we used an automatic parameterising algorithm (Haller et al., 2018) that efficiently disentangles the two features -the *slope* and the *offset* – that characterise the 1/*f* signal, and the three features – the *centre frequency*, *power* and *bandwidth* – that characterise the oscillatory components. Since the automatic parameterising algorithm does not impose band boundaries, this method allows for the assessment of group and individual differences in the *centre frequency*, *power* and *bandwidth* of the oscillations, both in the broadband spectra and in pre-defined narrow bands. With regard to possible age-associated differences in 1/*f* signal, there exists only one developmental fMRI/EEG study so far, which showed that the 1/*f* slope was significantly flatter in 17 healthy adults compared to 21 full-term newborns (Fransson et al., 2013). Based on the limited developmental MEG/EEG evidence regarding the 1/*f* signal, we hypothesised that differences in the 1/*f* signal may account to a large extent for observed age-related power differences between children and adults, and that canonical frequency band analyses may confound some band specific power changes with 1/*f* signal shifts. In particular, we predicted that the 1/*f* slope would be flatter, and the offset would be smaller in adults as compared to children (Fransson et al., 2013).

## METHODS

### Participants and Ethics Statement

This study included 52 human participants (28 children and 24 adults), namely healthy controls that had been recruited in a larg project on stuttering. Data from 4 children were excluded from the analysis due to excessive head movement (> 5 mm), incidental system noise or signs of drowsiness throughout the recording. Drowsiness was monitored online through a video-camera so that any affected data would be removed from further analysis. Child participants were accompanied by an experienced researcher who sat with them during the whole session to make sure they remained comfortable, and who monitored and encouraged their compliance. The final sample consisted of 24 children (8.0 ± 2.5 years, 17 males) and 24 adults (40.6 ± 17.4 years, 16 males).

Written informed consent was obtained from the adult participants and from the parents/guardians of the children prior to the experiment. No participant reported personal or family history of neurological disease or psychological impairment and none were taking medication that could affect MEG recordings at the time of participation. All participants were remunerated $AUD 40 for their participation. The experimental procedures were approved by the Human Participants Ethics Committee at Macquarie University.

### MEG Data Acquisition

Resting-state MEG data of 300 seconds were acquired for child and adult participants using two separate whole-head gradiometer MEG systems. Child data were acquired using a paediatric 125-channel whole-head gradiometer MEG system (Model PQ1064R-N2m, Kanazawa Institute of Technology/KIT, Kanazawa, Japan). Adult data were acquired using a 160-channel whole-head gradiometer MEG system (Model PQ1160RN2, KIT, Kanazawa, Japan). Use of the paediatric MEG system to overcome critical limiting factors for MEG experimentation on children below the ages of five to six years (Irimia et al., 2014), including a much smaller head size and overall structure in children (the smaller crown to shoulder distance prevents the full insertion of the head into the adult helmet), has been demonstrated previously (Sowman et al., 2014; He, Brock, et al., 2015; He, Garrido, et al., 2015; Etchell et al., 2016).

The gradiometers of both systems have 50 mm baseline and 15.5 mm diameter coils positioned in a glass fibre reinforced plastic cryostat for measurement of the normal component of the magnetic field from the human brain (Kado et al., 1999). In both systems, neighbouring channels are 38 mm apart and 20 mm from the outer dewar surface. These factors ensure that the signals obtained by the two MEG systems are equivalent. The 125-channel dewar was designed to fit a maximum head circumference of 53.4 cm, accommodating more than 90% of heads of 5-year-olds (see Johnson et al., 2010 for details). Both systems were situated within the same magnetically shielded room within the KIT-Macquarie Brain Research Laboratory (https://www.mq.edu.au/research/research-centres-groups-and-facilities/healthy-people/facilities/meg), and therefore environmental noise was comparable.

During MEG data acquisition, participants were asked to remain relaxed, awake and with their eyes fixed on a white cross at the centre of a black 36 cm (width) x 24 cm (length) rectangular image with 4 x 4 degrees of visual angle. Visual display was presented on a back-projected screen mounted approximately 140 cm above the participant using video projectors situated outside the magnetically shielded room (child MEG projector: Sharp Notevision Model PG10S, Osaka, Japan; Adult MEG projector: InFocus Model IN5108, Portland, USA). An overview of the child-friendly experimental protocol can be found in the video article (Rapaport et al., 2019).

### MEG Data Processing

An overview of the processing pipeline is illustrated in Figure 1. MEG data were acquired at a sampling frequency of 1000 Hz, using a hardware bandpass filter of 0.03-200 Hz. The continuous raw MEG data were filtered off-line from 0.5 to 100 Hz using bi-directional IIR Butterworth filters with DC removal and segmented into epochs of 4096 samples (= 4.096 seconds). The data were visually inspected by WH, and epochs that contained oculographic, myographic, and system/environmental artefacts (e.g., squid jumps) were removed. The first and last epochs were also excluded from the analysis. A mean of 23.8 ± 3.02 artefact-free epochs of 4.096 s data in children (15-28 epochs) and 40.0 ± 0.02 artefact-free epochs in adults (39-40 epochs) were selected for subsequent source modelling. There were age-related differences in the number of clean trials between children and adults (*t* (46) = 26.31, *p* < 0.01; two-sample t-test using the ttest2 function in MATLAB, version R2017b), as expected and was inevitable due to the fact that more trials were removed from younger participants because of movement. However, it has been shown in previous simulations that beamformer performance plateaus before the lower limit of ∼80 seconds of data that was used for our analysis (20 epochs of 4.096 seconds; Brookes et al., 2008).

**Figure 1.**
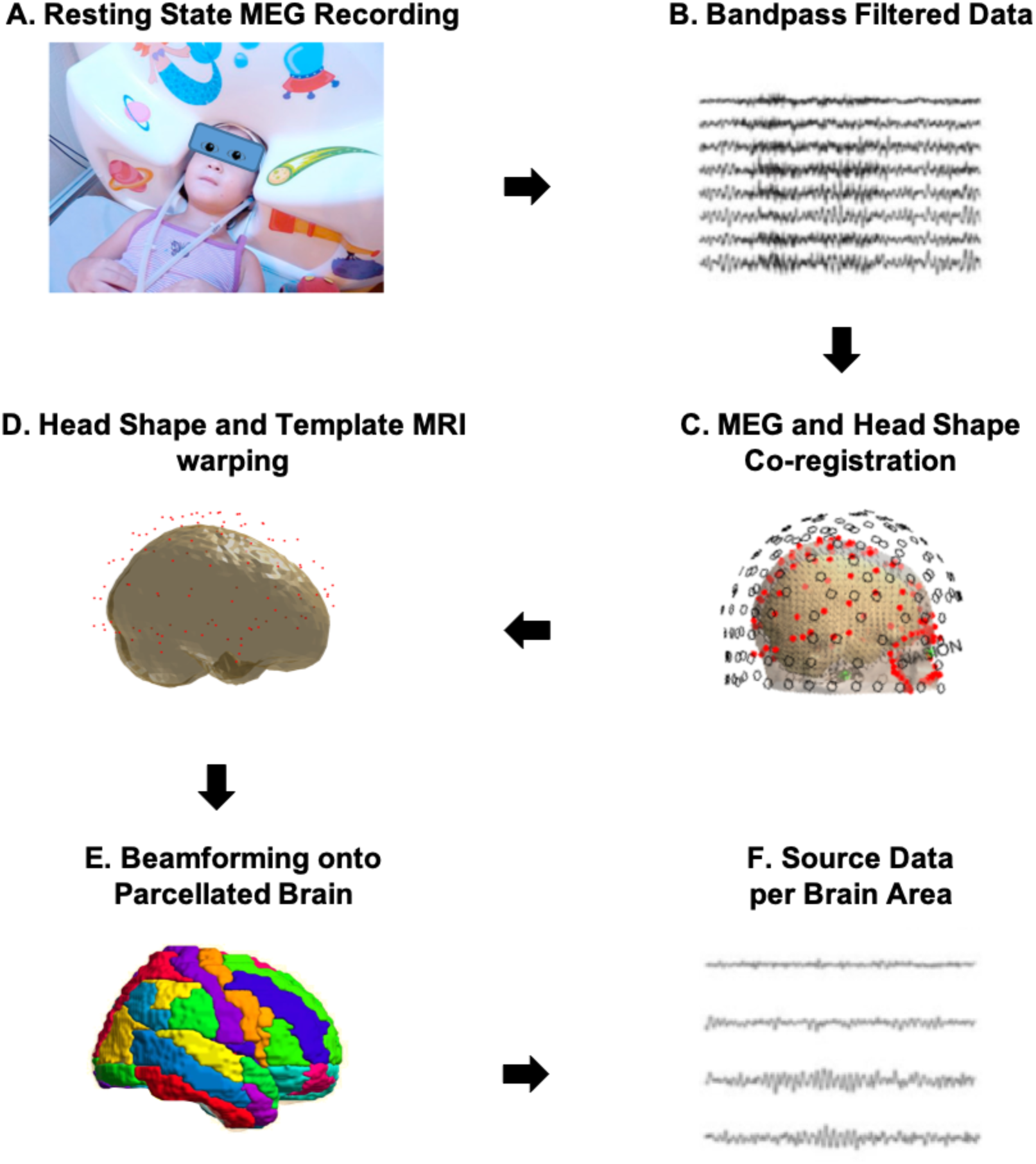
Schematic overview of the processing pipeline. The eyes-open resting state MEG were recorded (A) and bandpass filtered (B) before being co-registered to individual head shape (C) and template MRI (D). Following this, data were epoched and beamformed to parcels of the AAL atlas (E). Power spectral density was then estimated from individual cortical sources by Welch’s method (F).

MEG sensor data were then projected onto a parcellated cortical surface using an atlas-based beamforming approach (Hillebrand et al., 2012; Hillebrand et al., 2016), providing source activities at each centroid of the automated anatomical labelling (AAL) atlas (Tzourio-Mazoyer et al., 2002). Firstly, the geometry of each participant’s scalp was reconstructed from a “surrogate” MRI, where the Montreal Neurological Institute (MNI) template T1 structural brain image was warped to each participant’s digitized head shape with an iterative closest point algorithm implemented in *BrainWave* (Cheyne et al., 2014). Secondly, a multi-sphere volume conductor model was calculated using the outline of the scalp from this co-registered data in MRIViewer of the CTF MEG5 software (VSM MedTech Systems Inc., Coquitlam BC, Canada; Version 5.0.2). Thirdly, the broadband (0.5-48 Hz) data covariance matrix was calculated from all selected epochs, and a unity noise covariance was used. Lastly, the data covariance, the unity noise covariance, together with an equivalent current dipole source model and the multi-sphere volume conductor model, were combined to reconstruct beamformer weights for the parcels’ centroids using Synthetic Aperture Magnetometry (SAM, Robinson, 1999). Subsequently, the broadband MEG sensor data were projected through the normalised beamformer weights in order to obtain traces of neuronal activity in the cortical space (Cheyne et al., 2007).

To counteract trial imbalance between groups, we chose for each individual the first 15 artefact-free epochs from each of the 80 AAL source regions of interest (80 Regions of Interest/ROIs; 78 cortical and bilateral hippocampal) for the subsequent estimation of power spectral density.

### Spectral Analysis

#### Conventional Analysis in *a priori* Defined Frequency Bands

Power spectral density (PSD) was estimated for each participant, ROI and artefact-free epochs separately using Welch’s method (Welch, 1967) implemented in MATLAB 2017b, with 50% overlap and a Hamming window of 3s (resulting in a spectral resolution of 0.24 Hz). A single PSD for each participant was obtained by averaging the PSDs across all epochs and ROIs.

A conventional spectral analysis was carried out by calculating the absolute power from the raw power spectrum in five canonical frequency bands (delta: 1–4 Hz, theta: 4–8 Hz, alpha: 8–13 Hz, beta: 13-30 Hz, and low gamma: 30–48 Hz). The frequency at which each participant reached the peak amplitude was calculated within the 5-13 Hz band for children and the 8-13 Hz band for adults using an automated local maxima algorithm (MATLAB function findpeaks). The lower frequency boundary was used for children in order to account for reduced alpha peak frequencies in young children when compared to adults (Klimesch, 1999; Bathelt et al., 2013; Mierau et al., 2016).

#### Parameterising the Power Spectrum with no *a priori* Defined Frequency Bands

The PSDs, calculated by Welch’s method, were also submitted to the FOOOF v0.1.3 parameterisation model – an open source Python package (https://github.com/fooof-tools/fooof/; in Python v3.7.0) - for automatic separation of periodic and aperiodic components of neural power spectra (Haller et al., 2018). Briefly, the model considers the PSDs as a linear sum of aperiodic “background” neural signal (1/*f* signal) and oscillations, or peaks represented by Gaussian functions in the PSD, above the 1/*f* signal level.

The power spectrum % is then modelled as:

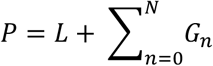

with *L* the aperiodic “background” signal and *G* the oscillations, modelled as *N* Gaussians. The aperiodic signal is fitted, after which the aperiodic fit is subtracted from the power spectrum, creating a flattened (or aperiodic-adjusted) spectrum, wherein peaks were iteratively fitted by Gaussians modelled as:

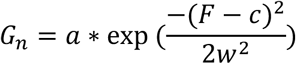

where *a* is the amplitude, *c* is the centre frequency, *w* is the bandwidth of the Gaussian *G*, and *F* is the vector of input frequencies.

Subsequently, a peak-removed power spectrum is calculated by subtracting all fitted Gaussians from the original power spectrum. Finally, an aperiodic signal is re-estimated from this peak-removed power spectrum, representing the cortical 1/*f* background signal. Both the initial and final fit of the aperiodic component are fit as:

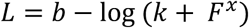

where *b* is the broadband offset, *x* is the slope, *k* and is the “knee” parameter, which indicates where the “bend” occurs in the 1/*f* component. In the current case of non-invasive MEG recordings, no knee was expected across the frequency range studied (Miller et al., 2009).

The FOOOF model was fitted across the frequency range of 1 to 48 Hz in fixed (no spectral knee) mode (*peak_width_limits* = [0.5, 12], *max_n_peaks* = 10, *min_peak_amplitude* = 0, and *peak_threshold* = 2, *aperiodic_mode* = ‘fixed’). Goodness-of-fit of the FOOOF model is returned in terms of the *R*^2^ of the fit.

Two parameters - the **Slope x** and the **Offset b** – defining the aperiodic 1/*f* background signal, and three parameters – the **Centre Frequency *c***, **Power *a*** and **Bandwidth *w*** – characterising the oscillations were returned by the FOOOF parameterisation, and entered permutation statistical comparisons between the two age groups.

In addition, the final aperiodic signal returned from the FOOOF model was subtracted from the raw power spectrum, resulting in a flattened spectrum from which the absolute alpha and beta band power was estimated again using the conventional spectral approach.

### Statistical Analysis

Statistical analyses were performed using permutation testing as implemented in the Resampling Statistical Toolkit for MATLAB 2017b (https://au.mathworks.com/matlabcentral/fileexchange/27960-resampling-statistical-toolkit). We used 50,000 permutations of group membership to empirically approximate the distribution for the null hypothesis (i.e., no difference between groups) for each contrast. For each permutation, t-values were derived for a contrast of interest, and any t-values for the original contrast that exceeded the significance threshold of 0.05 for the t-distribution were deemed reliable.

In addition, Bayes Factors (alternative BF/null BF; Wagenmakers, 2007; Dienes, 2011) were estimated in JASP v0.9.2 (Quintana and Williams, 2018; https://jasp-stats.org/) to further quantify the effect size, and to facilitate the interpretation of evidence for or against the null hypothesis when comparing to the alternative hypothesis. For the alternative hypotheses of measures being larger in adults than children and vice versa, a “unit information prior” was assumed with a default Cauchy prior with a scale parameter of 0.707 (Jeffreys, 1998). Bayesian correlation analyses, which allows inferences on the absence of a correlation between variables to be made, were also conducted in JASP. For testing the correlations, a prespecified alternative hypothesis with a flat beta prior width of 1 centred around K = 0 was used for a null hypothesis (*r* = 0). An illustration of the effects of assigning a range of different prior distributions (i.e., a Bayes factor robustness check) was conducted for all Bayesian tests.

BFs were thresholded > 3, > 10, and >100 as substantial, strong, and very strong/decisive evidence in favour of the alternative hypothesis, and BF < 1/3 and < 1/10 for substantial and strong evidence for the null hypothesis (Raftery, 1995; Jeffreys, 1998). BFs that fell in-between 1/3 and 3 were taken as insufficient evidence for either hypothesis (Jeffreys, 1998; Dienes, 2014).

## RESULTS

### Raw Power Spectra With *a priori* Defined Frequency Bands

#### Decreases in Low- and Increases in High-Frequency Power

Figure 2A&B depicts the results of conventional spectral analyses, absolute power was computed in five *a priori* defined frequency bands (delta: 1–4 Hz, theta: 4–8 Hz, alpha: 8–13 Hz, beta: 13-30 Hz, and lower gamma: 30–48 Hz). The PSD in Figure 2A shows a tendency of the grand average power for adults to be lower across lower frequency bands, and higher in higher bands, compared to children. Permutation statistics identified significantly lower power in the delta (Figure 2B, *t*(46) = −6, *p* < 0.01) and theta (*t*(46) = −3.78, *p* < 0.01) bands, and higher power in the beta (*t*(46) = 2.74, *p* < 0.01) and gamma bands (*t*(46) = −2.46, *p* = 0.01) in adults compared to children. The right-corner panel in Figure2A shows that the peak frequency was significantly higher in adults (9.99 ± 1.30 Hz) than in children (7.58 ± 1.71 Hz, *t*(46) = 5.5, *p* < 0.01).

**Figure 2.**
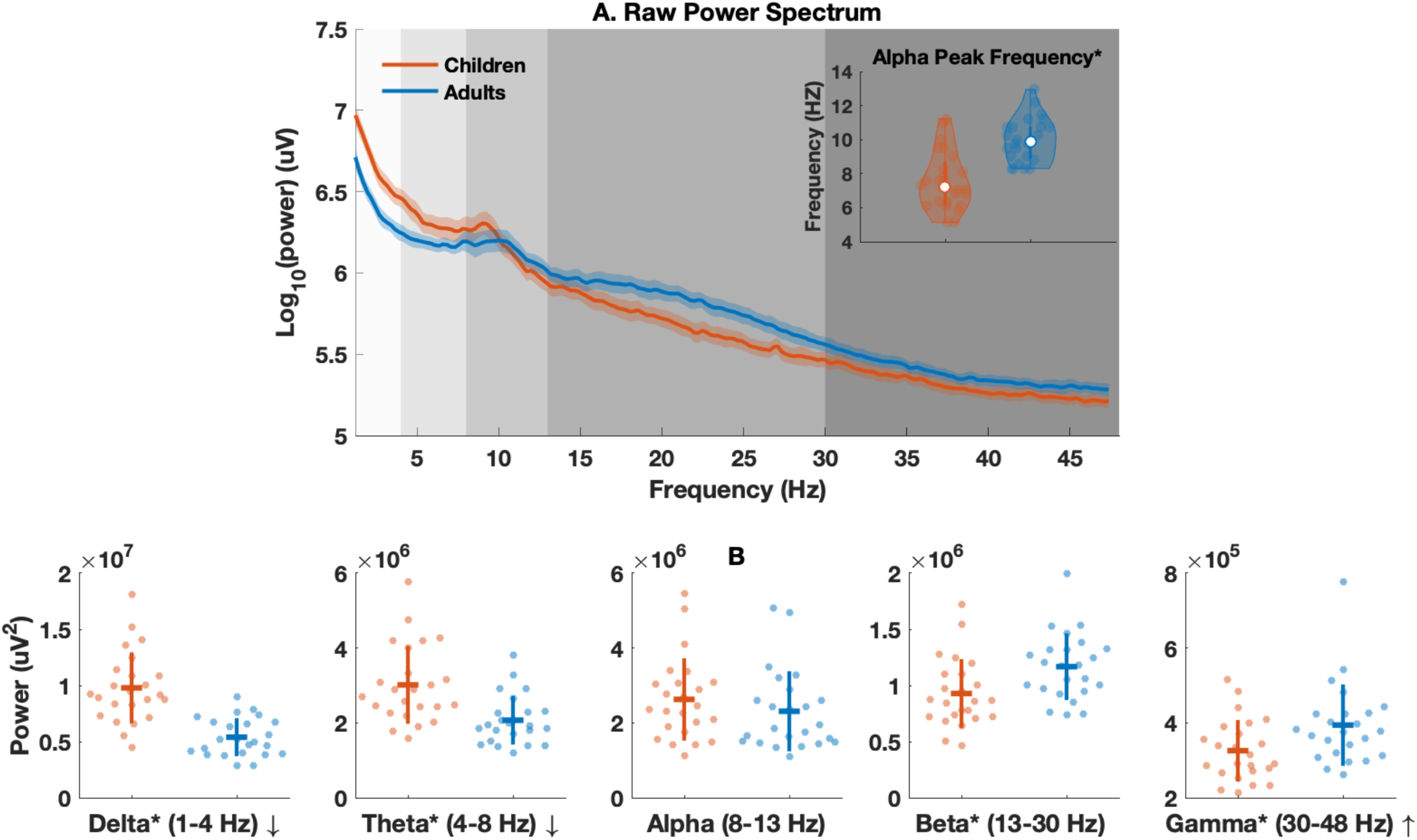
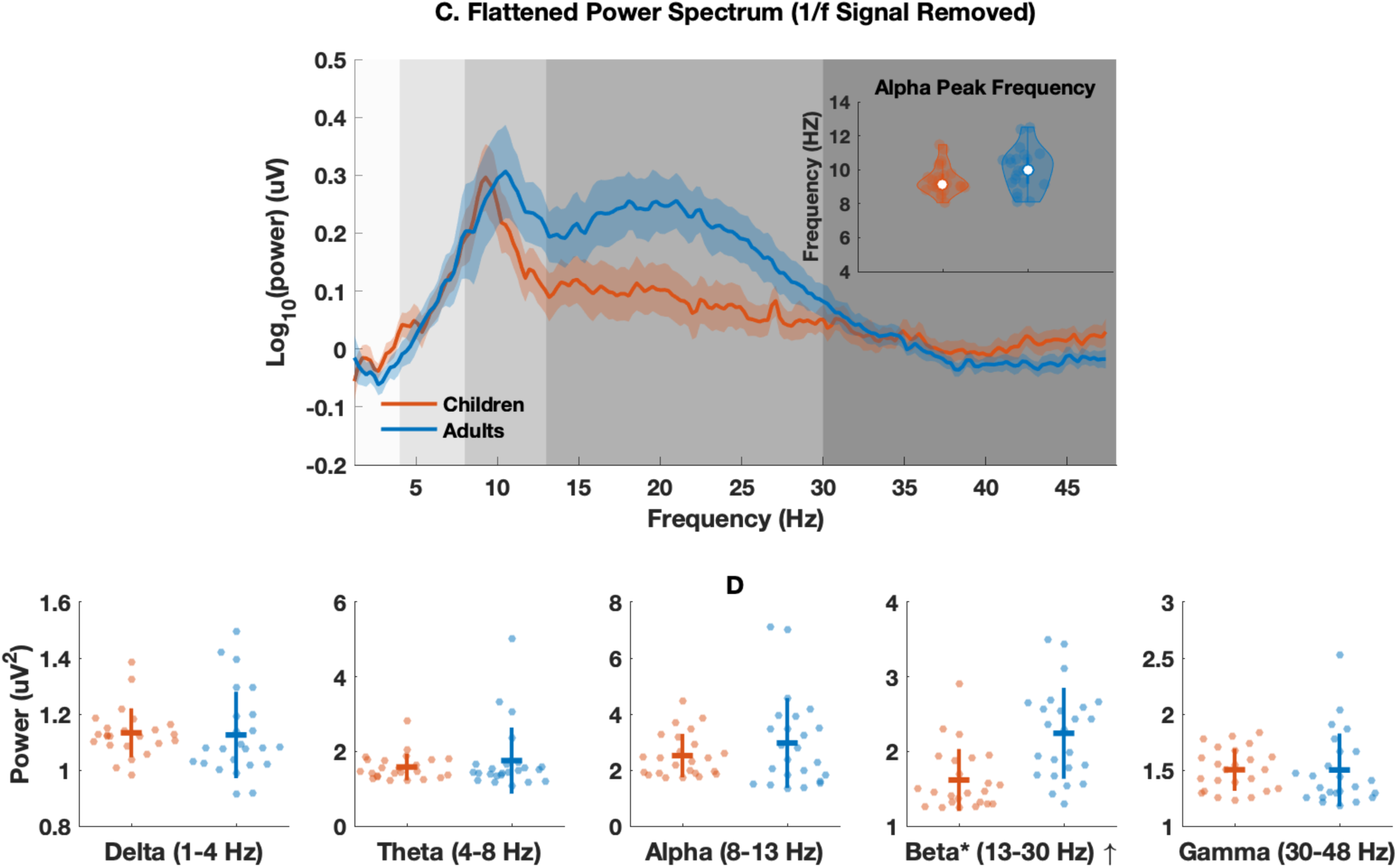
Power Spectral Density for Children (in red, N = 24) and Adults (in blue, N = 24). Grand average of the raw power spectrum (A) and flattened power spectrum (C) for children and adults, with 95% confidence intervals represented by shaded areas (Gaussian-distribution assumed). Violin plots of the peak frequencies identified in the raw power spectra for children (5-13 Hz) and adults (8-13 Hz) are shown in the top-right panels of Figure 2A&C. The white dots depict group mean values. Power (B) and aperiodic-adjusted power (D) in five *a priori* defined canonical frequency bands as colour coded regions with grey gradients. Each dot represents a child participant in read or an adult participant in blue. Horizontal lines indicate group mean and vertical lines standard deviations. * indicates group difference reaches statistical significance; ↑ and ↓ indicate significant increase and significant decrease in power, respectively.

In addition, the data were examined by estimating a Bayes factor (alternative BF/null BF), which indicates the fit of the data under the alternative hypothesis. Estimated BFs indicated decisive evidence for lower PSDs in the delta (BF > 100, 95% Confidence Interval = [−2.30, - 0.93]) and theta (BF = 119.80, 95% CI = [−1.58, −0.41) bands, higher PSDs in the beta (BF = 10.66, 95% CI = [0.16, 1.26]) and gamma (BF = 6.13, 95% CI = [0.10, 1.18]) bands, and higher peak frequency (BF > 100 95% CI = [0.86, 2.13]) in adults than children. Bayes Factors suggested that there was strong evidence for the absence of a group difference in alpha band power (BF = 0.16, 95% CI = [0.01, 0.44]).

### Parameterised Power Spectra Without *a priori* Defined Frequency Bands

#### Increase in 1/*f* Slope and Decrease in 1/*f* Offset

Figure 3A shows the aperiodic component of the grand average parameterised spectrum (i.e., peaks were removed from the power spectrum) for each age group. The aperiodic power spectrum was flatter in adults than in children. Permutation testing confirmed that the 1/*f* slope (Figure 3B) was significantly different (flatter) in adults compared to children (adults = −0.89 ± −0.12; children = −1.15 ± −0.09, *t*(46) = 8.59, *p* < 0.01), and the 1/*f* offset was smaller (adults = 6.78 ± 0.15; children = 7.11 ± 0.17, *t*(46) = -6.99, *p* < 0.01). Goodness-of-fit, as indexed by R^2^ of the modelling fit was 0.99 ± 0.01 for adults, and 0.99 ± 0.01 for children (*t*(46) = −1.05, *p* > 0.05), suggesting that a difference in the model fit was not the cause of observed differences in the 1/*f* signal between groups.

**Figure 3.**
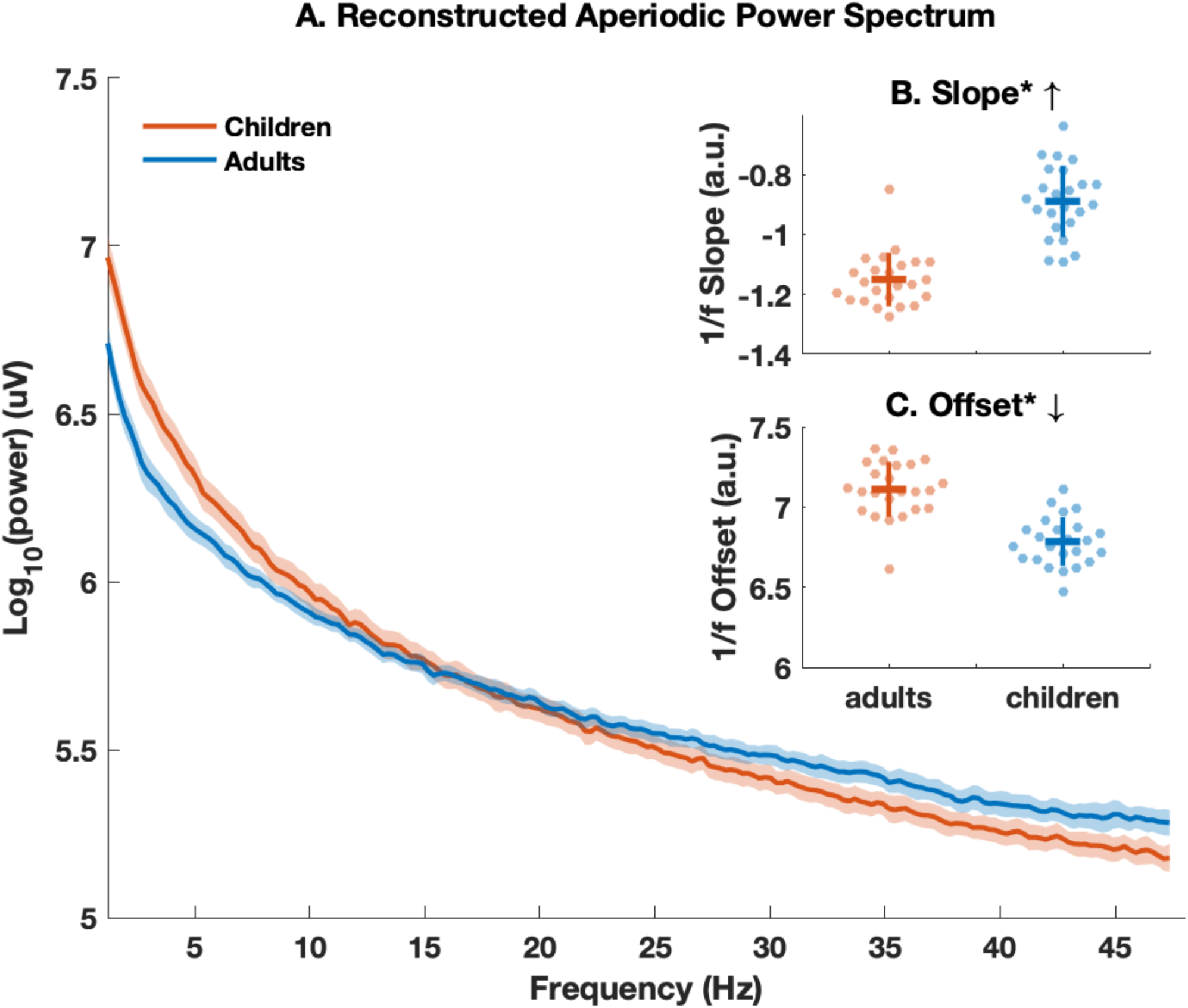
Non-oscillatory Power Spectrum (i.e., oscillatory components have been removed) for Children (in red, N = 24) and Adults (in blue, N = 24). A: Grand average of the log-transformed power spectrum of the aperiodic signal for children and adults, with 95% confidence intervals represented by shaded areas (Gaussian-distribution assumed). B&C: Slope and Offset of the aperiodic power spectrum for both groups. Horizontal lines indicate group mean and vertical lines standard deviations. * indicates group difference reaches statistical significance; ↑ and ↓ indicate significant increase and significant decrease in power, respectively.

Bayes Factors indicated decisive evidence for flatter/more positive 1/*f* slope (BF > 100, 95% CI = [1.66, 3.15]) and lower 1/*f* offset (BF > 100, 95% CI = [−2.62, −1.20]) in adults compared to children.

#### Correlation between Age and the Aperiodic **1/*f*** Component

Bayes Factors revealed a decisive negative correlation between age and 1/*f* offset (*r* = - 0.71, BF > 100, 95% CI = [−0.85, −0.39]), and a positive correlation between age and 1/*f* slope (*r* = 0.62, BF = 35.88, 95% CI = [0.26, 0.8]) in children. This trend became anecdotal in adults (offset: *r* = −0.4, BF = 1.56, 95% CI = [−0.67, 0.01]; slope: *r* = −0.5, BF = 3.7, 95% CI = [0.14, 0.75]).

#### Power and Bandwidth Increase for Beta Power Peaks

Figure 4 shows the group comparisons for the periodic components - **Centre Frequency**, **Power**, and **Bandwidth** - of the peak oscillation (i.e., the highest power peak across all frequencies in FOOOF model). Figure 4A demonstrates that 95.83% of children and 58.33% of adults exhibited oscillatory peaks that fall within the alpha range, whereas only 4.17% children but 41.63% adults had a peak oscillation in the beta band. This suggests that more adults had oscillatory peaks outside of the canonical 8-13 Hz alpha range. Permutation comparisons showed that for the peak oscillation the centre frequency was significantly higher in adults compared to children (Figure 4B; adults = 13.52 ± 4.86, children = 10.25 ± 3.97, *t*(46) = 2.55, *p* = 0.01; BF = 7.36, 95% CI = [0.13, 1.21]), and the bandwidth was significantly larger in adults than in children (Figure 4D; adults = 3.63 ± 3.35, children = 1.80 ± 1.13, *t*(46) = 2.53, *p* = 0.01; BF = 7.07, 95% CI = [0.12, 1.2]).

**Figure 4.**
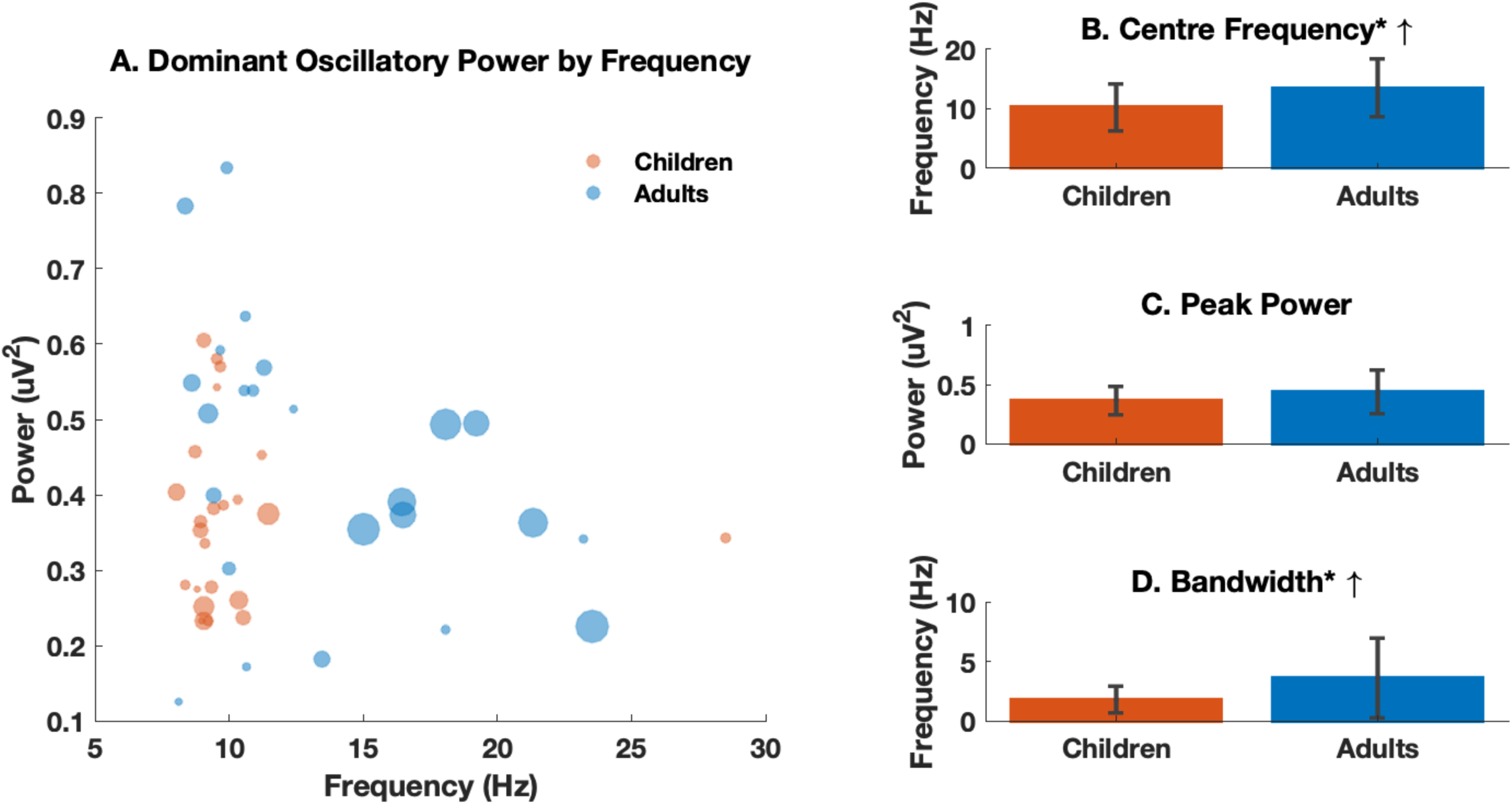
Dominant Oscillations in the Parameterised Power Spectrum for Children (in red, N = 24) and Adults (in blue, N = 24). A: The power of the dominant oscillation (i.e., oscillation with the maximum power across the broadband power spectrum) by frequency for each individual, with the size of the circle representing the bandwidth of the dominant oscillation. B-D: Statistical comparisons of the centre frequency, power and the bandwidth of the dominant oscillation. Error bars represent standard error of the group means. * indicates group difference reaches statistical significance; ↑ and ↓ indicate significant increase and significant decrease in power, respectively.

In order to make a valid comparison between parameterised oscillatory components and canonical narrowband analysis, the centre frequency, power, and bandwidth of the highest oscillatory component were also extracted independently for the alpha and beta bands from all individuals (Figure 5A). Figure 5B&E shows Gaussian curves obtained from the individual oscillatory component values of the FOOOF model for the alpha and beta bands. There is significant individual variability observable for both bands, which is further quantified in Figures 5C, D, F and G. The aperiodic-adjusted beta peak oscillation (but not alpha) was significantly higher in power (Figure 5F; adults = 0.31 ± 0.11, children = 0.18 ± 0.09, *t*(46) = 4.37, *p* < 0.01; BF > 100, 95% CI = [0.51, 1.77]) and larger in bandwidth (Figure 5G; adults = 4.87 ± 3.44, children = 2.39 ± 2.58, *t*(46) = 2.83, *p* = 0.01; BF = 13.01, 95% CI = [0.15, 1.29]) in adults compared to children.

**Figure 5.**
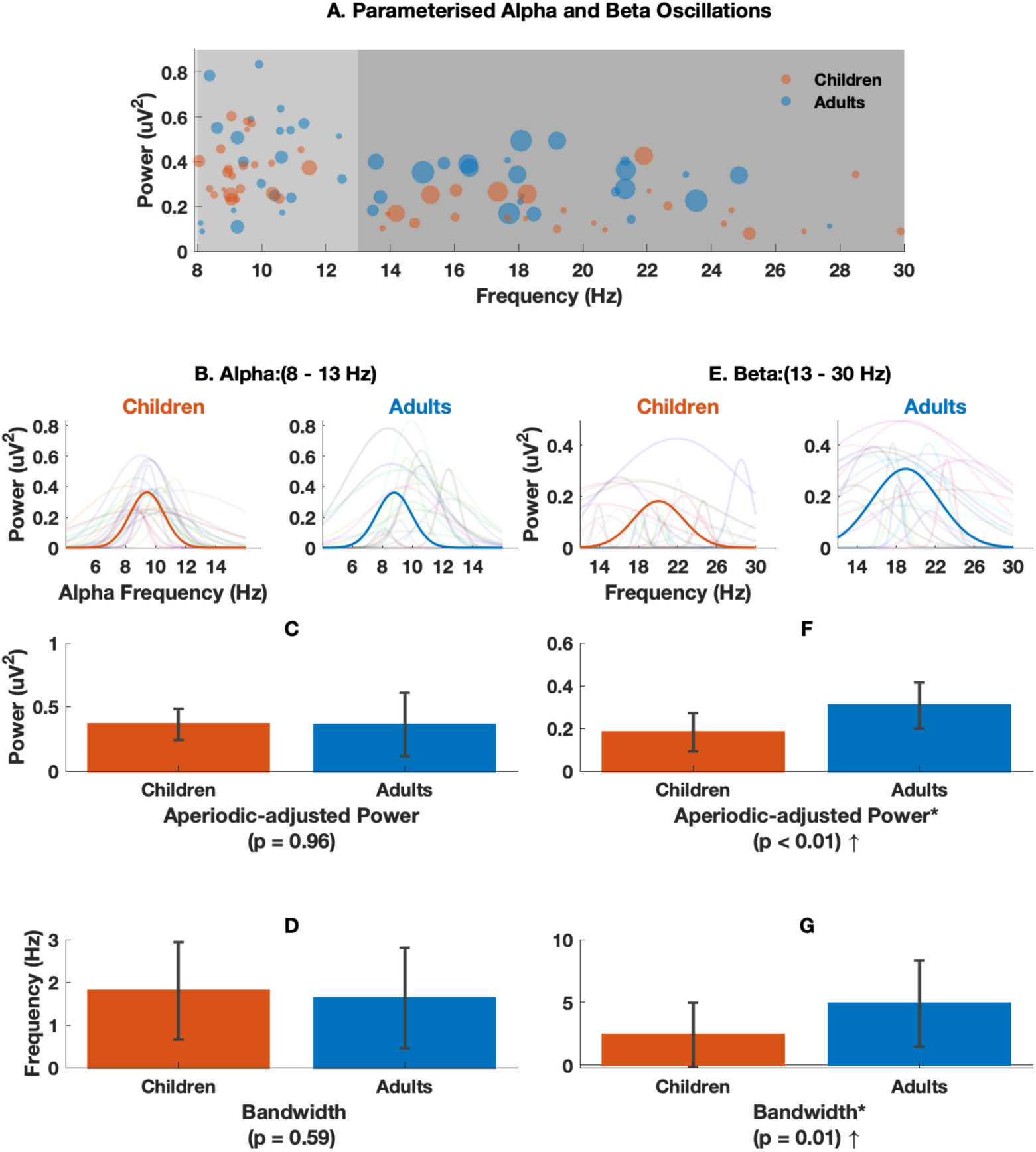
Alpha and Beta Band Parameterisation of the Power Spectrum for Children (in red, N = 24) and Adults (in blue, N = 24). Two-band models of the power spectrum (A) – alpha (red, 8–13 Hz) and beta (blue, 13-30 Hz). B&E show the maximum oscillatory power and the frequency at which this occurs for each individual. Statistical comparisons of the power (C&F) and the bandwidth (D&G) of the oscillations in the alpha and beta bands. Error bars represent standard error of the group means. frequency, power and the bandwidth of the peak oscillation components. Error bars represent standard error of the group means. * indicates group difference reaches statistical significance; ↑ indicates significant increase in power.

#### Correlation between Age and Peak Oscillatory Components

Bayesian analysis only identified a very strong positive correlation between age and bandwidth in all participants (*r(46)* = 0.47, BF = 43.08, 95% CI = [0.2, 0.65]), but no such evidence for the individual age groups (adults: *r* = 0.4, BF = 1.47, 95% CI = [−0.11, 0.67]; children: *r(46)* = 0.26, BF = 0.51, 95% CI = [−0.16, 0.57]).

In addition, when analysing the periodic components in the beta and alpha bands, Bayes Factors revealed a very strong positive correlation between age and power in the beta band in all participants (*r* = 0.5, BF = 43.08, 95% CI = [0.2, 0.65]), but substantial evidence for the absence of such a correlation in individual age groups (adults: *r* = 0.08, BF = 0.27, 95% CI = [−0.31, 0.44]; children: *r* = 0.07, BF = 0.27, 95% CI = [−0.32, 0.43]). In children a strong positive correlation was found between age and centre frequency in the alpha band (*r* = 0.61, BF = 27.14, 95% CI = [0.24, 0.8]).

#### Beta-specific Power Increase in the Flattened Parameterised Power Spectra

Figure 2C&D displays the results of conventional spectral analysis applied to the flattened PSD, where the aperiodic signal was removed from the power spectrum, thereby leaving only the oscillatory components. Interestingly, unlike the multiple band-specific power differences found in the raw power spectra (Figure 2B), the permutation analysis only identified significantly increased beta power (Figure 2D, *t*(46) = 4.15, < 0.01) when comparing adults and children. The same frequency ranges (children: 5-13 Hz; adults: 8-13 Hz) as for the conventional analyses were applied when estimating peak frequencies in FOOOF. The right-corner panel in Figure2B shows that there was no significant difference in the peak frequency between adults (8.78 ± 3.58 Hz) and children (9.42 ± 0.84 Hz, *t*(46) = −0.85, *p* = 0.4) in the flattened spectra.

Bayes Factors revealed decisive evidence for larger aperiodic-adjusted power in the beta band (BF > 100, 95% CI = [0.45, 1.72]) in adults compared to children, insufficient evidence for any group difference in the delta and alpha bands (delta: BF = 0.35, 95% CI = [−0.64, −0.01]; alpha: BF = 0.92, 95% CI = [0.02, 0.85]), and strong evidence for the absent group differences in the theta and gamma bands (theta: BF = 0.17, 95% CI = [−0.46, −0.006]; gamma: BF = 0.28, 95% CI = [0.01, 0.59]). Bayes Factors further suggested substantial evidence for absence of group difference for alpha peak frequency (BF = 0.17, 95% CI = [0.01, 0.45]).

#### Positive Correlation between Aperiodic-adjusted Beta Power and Aperiodic 1/*f* Component

Based on the significant group differences in the aperiodic-adjusted power in the beta band and 1/*f* signal identified in the parametrised spectra, the presence of correlations between the beta power in the flattened spectra and the parameters of the 1/*f* signal was assessed. Bayes Factors revealed a very strong correlation between the narrowband aperiodic-adjusted beta power in the flattened spectral and 1/*f* Slope (*r* = 0.5, BF = 94.36, 95% CI = [0.24, 0.67]), and 1/*f* Offset (*r* = −0.49, BF = 66.53, 95% CI = [−0.66, −0.22]), across all participants but not in individual age groups.

## DISCUSSION

We investigated developmental changes in the aperiodic 1/*f* and the oscillatory components of MEG brain signals. The findings of our conventional analyses of narrowband power and peak frequency were remarkably similar to previous studies (Marcuse et al., 2008; Boersma et al., 2011; Cragg et al., 2011; Miskovic et al., 2015; Gomez et al., 2017; Rodriguez-Martinez et al., 2017). However, these results turned out to be mostly non-significant (except in the beta band) once the aperiodic 1/*f* signal and the oscillations were disentangled, and once the narrowband power was assessed in the flattened power spectrum (that is after the 1/*f* component had been removed as shown in Figure 2). In these non-conventional analyses, we observed distinct (complementary) developmental profiles for the 1/*f* signal and oscillatory power, with compelling evidence of flatter 1/*f* signals and increased beta oscillations in the adults, as compared to the children. Moreover, the strong correlation between the 1/*f* signal and beta oscillatory power suggested a co-maturation of the two neural phenomena during child development.

### Conventional versus Parameterised Power Spectral Analyses

Previous developmental studies have shown power decreases in lower frequency bands (Puligheddu et al., 2005; Gomez et al., 2013; Schafer et al., 2014), and increases in alpha peak frequency throughout childhood (Marcuse et al., 2008; Boersma et al., 2011; Cragg et al., 2011; Smit, Boomsma, et al., 2012; Miskovic et al., 2015; Gomez et al., 2017; Rodriguez-Martinez et al., 2017). By applying conventional methods, we were able to replicate these findings. However, none of these findings remained significant once careful adjudication between aperiodic 1/*f* and oscillatory components was carried out using power spectra parameterisation (Haller et al., 2018).

Interestingly, the peak frequency in the flattened spectra was found to correlate positively with age in the child participants only, but the peak frequency showed no differences between age groups (Figure 2B), and neither did the aperiodic-adjusted power or frequency bandwidth in the alpha band (Figure 5C&D). These findings contrast with significant age differences found in the alpha band when not accounting for the aperiodic 1/*f* signal, and suggest that the magnitude of these changes could be heavily conflated by the aperiodic 1/*f* signal in the raw power spectrum. This is further supported by the significant correlation that was identified between the 1/*f* signal (both offset and slope) and age, only in the child participants. These findings invite the conclusion that developmentally-related power decreases in lower frequency bands and peak frequency increases that have been revealed by standard methods are at least partially driven by the flattening of the 1/*f* component in the broadband spectrum (Voytek and Knight, 2015).

### Flatter Slope Indicates Increase in Neuronal Noise

The slope of the 1/*f* signal was found to be flatter, or less negative, in adults than in children. According to the “Wiener-Khinchin theorem”, the power spectrum is equivalent to the Fourier transform of the autocovariance function (He, 2014). Therefore, a flatter slope in the frequency domain indicates a shorter/weaker autocorrelation in the time domain. Interestingly, the reduction of the autocorrelation in human brain activity has been found to correlate with the increasing demands for more efficient online information processing during cognitive load in working memory tasks (He, 2014; Voytek et al., 2015). Indeed, a reduced temporal integration span in brain activity would be expected to reflect the need for enhanced integration of information during brain development.

There is also evidence that neuronal spiking statistics are relevant to the 1/*f* slope (Voytek and Knight, 2015; Gao, 2016), in that the slope of the aggregated local field potential becomes flatter when a large number of spikes occur asynchronously (Usher et al., 1995; Pozzorini et al., 2013; Voytek and Knight, 2015). This decoupling of neuronal population spiking from an oscillatory regime, which has been broadly defined as “noise”, can be driven by increases in the ratio between local excitation/inhibition (Cremer and Zeef, 1987; McIntosh et al., 2010; Hong and Rebec, 2012; He, 2014; Voytek et al., 2015). In fact, the inhibitory regulation of the synchronisation between pyramidal neurons from the GABAergic neurons has been found to undergo prolonged changes into adolescence (Hashimoto et al., 2009).

The increase of neuronal noise during brain development has also been reported in studies adopting a simplistic method that uses the variability of brain signals as a proxy for neuronal noise. Such efforts have consistently identified increasing variability of both spontaneous and evoked brain activity to correlate with more stable and accurate behaviour during development (McIntosh et al., 2010; Fransson et al., 2013). Of direct relevance to our findings, a cross-sectional EEG study comparing children aged 8-15 years with young adults reported that the variability of evoked face responses increased with age. This variability turned out to be positively correlated with intrasubject response accuracy but negatively correlated with reaction-time variability (McIntosh et al., 2008). This increase in neuronal noise in the developing brain appeared to be more global, as compared to a more local noise increase with aging (McIntosh et al., 2010). Speculatively, neuronal noise during childhood shapes the brain from a deterministic system into one that is more stochastic and adaptive to an uncertain environment (Knill and Pouget, 2004; Stein et al., 2005). Brain noise in the aging population continues to increase, but the pattern of changes can be more focal and is often in parallel with diminished neuroplasticity (Li et al., 2006; Garrett et al., 2010). Such speculation regarding the network level gains support from our recent analysis of brain network topology in the same dataset, where pervasive decreases in connectedness were revealed (i.e., nodal centrality) in most cortical regions, with increasing global network segregation (He et al., 2019). The same approach, however, has also identified quite focal changes in hub regions as the driving force of large-scale network abnormalities in various neurological diseases (Stam et al., 2009; Crossley et al., 2014; DeSalvo et al., 2014; Tewarie et al., 2014; Yu et al., 2017).

### Smaller Offset Indicates Reduction in Broadband Power

Accumulating evidence suggests that the offset of the 1/*f* signal reflects the broadband power, which in turn is associated with the aggregated spiking activity of the underlying neuronal populations (Manning et al., 2009; Miller et al., 2009; Miller et al., 2014). We found a significant reduction in the offset of the 1/*f* signal for adults as compared to children, which could indicate a reduction in broadband power. Broadband power reductions have been reported consistently in developmental MEG/EEG studies (Miskovic et al., 2015; Gomez et al., 2017; Rodriguez-Martinez et al., 2017). This finding also corroborates a previous report on the developmental parallelism between the reduction in spectral power and cortical thickness (Whitford et al., 2007). It is understood that brain development begins with neuronal proliferation and synaptogenesis, followed by synaptic pruning during which synapses are selectively eliminated (Marsh et al., 2008). Computational work has shown that MEG signals represent spikes and the envelope of membrane de-/hyper-polarisation, predominantly from pyramidal neurons (Murakami and Okada, 2006). Therefore, one possible mechanism underlying the observed offset decreases is “regressive” cortical organisation due to the loss of grey matter (Giedd et al., 1999; Sowell et al., 2003).

We note here that our interpretation of the reduced 1/*f* offset needs to be taken with caution. This is because slope and offset of the 1/*f* signal are highly correlated (Haller et al., 2018), and thus any change in slope would be accompanied by a change in the offset, regardless of the offset shifts caused by broadband power reductions. Future studies are needed to clarify the extent to which the observed offset differences between age groups were caused over and above those expected from slope shifts.

### Co-maturing Beta Oscillations during Childhood

Analyses of oscillatory power revealed larger beta power in adults, as compared to children, and the difference remained significant once the 1/*f* signal was removed from the broadband power spectrum (Figures 6). The automatic parameterisation algorithm also revealed age-group differences in both peak oscillatory power and bandwidth in the beta band (Figures 5). Our findings add to previous developmental evidence for an increase in beta power using resting-state MEG/EEG (Puligheddu et al., 2005; Gomez et al., 2013; Schafer et al., 2014; Khan et al., 2018), and concurrent EEG-fMRI (Luchinger et al., 2011). These findings are also in accord with our recent longitudinal MEG study of motor development in children, which demonstrated linear increases in amplitude and mean frequency of beta (but not gamma) in movement (Johnson et al., 2019). Indeed, the beta band has been found to be heavily engaged in a wide range of processes such as cognitive control (Buschman and Miller, 2014), which develop well into adolescence (Luna et al., 2015) and changes with aging (Xifra-Porxas et al., 2019).

It is also worth noting that, in contrast to functional resonance imaging (fMRI), the MEG signal has a more direct relationship with neuronal activity and, compared to EEG signal, is less contaminated by age-related changes of structures external to the brain, such as the decreasing electrical conductivity of the human skull with age (Hamalainen et al., 1993; Hoekema et al., 2003; Baillet, 2017). Therefore, the increased absolute beta power detected by MEG reinforces the neuronal origin of the maturational power change reported by EEG and fMRI (Gasser et al., 1988; Marcuse et al., 2008; Boersma et al., 2011; Cragg et al., 2011; Smit, Boomsma, et al., 2012; Bathelt et al., 2013; Fransson et al., 2013; Miskovic et al., 2015; Gomez et al., 2017; Rodriguez-Martinez et al., 2017; Vandenbosch et al., 2019). In future work, we plan to determine if the current findings can be replicated in larger (longitudinal) data sets, and we will attempt to establish a more fine-tuned cortical representation of the 1/*f* and oscillatory signals from power spectra parameterisation. We encourage others to collect or reanalyse their resting-state data using the techniques we have employed (e.g., https://github.com/fooof-tools/fooof/), as well as other open source tools, to help establish norms of neuronal oscillations and aperiodic signals that are characteristic of healthy brain development.

### Limitations

There are some limitations to the present study. First, although our study provides valuable insights into age-group differences in key measures of neuronal activity, the cross-sectional nature of the study did not permit us to observe longitudinal changes at an individual level. A large longitudinal sample with balanced gender would be expected to replicate the current findings. Second, we assessed age-related differences based on 15 clean trials per participant. This was done to counteract differences in the number of clean trials between the adult and child participants. Data from the younger participants were prone to a greater number of artefacts, such as in-scanner motion. Future study should make use of a more efficient data acquisition approach (e.g., using calming video clips (Vanderwal et al., 2015)) to obtain larger data samples with improved quality in young children (Rapaport et al., 2019).

### Conclusions

By carefully modelling the power spectra of source-space electrophysiological data in adults and in children aged 4 to 12 years, the present study demonstrated strong evidence of correlated increases in the 1/*f* signal and beta power during child brain development. Our findings suggest that the reported power decreases in low-frequencies and the reported power increases in high-frequencies (other than the beta band) in many previous studies could have been caused by a flattening of the 1/*f* signal, instead of an authentic power change in the oscillations. In addition, our findings provide empirical support for the theory that neuronal noise increases with normal brain maturation (McIntosh et al., 2010). This change may shape the brain into a more stochastic system with balanced neuronal excitation/inhibition, greater complexity, and greater capacity for information processing (Knill and Pouget, 2004; Stein et al., 2005). Overall, the findings of the present study suggest that co-increasing beta oscillations and aperiodic brain signals accompany brain maturation during childhood.

## ACKNOWLEDGEMENTS

We thank all participants for their participation. We also thank Douglas Cheyne and Cecilia Jobst for their assistance in the preliminary data analysis, and Craig Richardson for his technical support in establishing inter-institutional collaborative data analysis platform. Finally, we acknowledge the collaboration of Kanazawa Institute of Technology in establishing the KIT-Macquarie MEG laboratory. This work was supported by the Australian Research Council (ARC) Centre of Excellence in Cognition and its Disorders (grant number CE110001021, http://www.ccd.edu.au), and the ARC Discovery Project (DP170103148). Wei He was supported by the Macquarie University Research Fellowship (#9201501199). Paul F. Sowman was supported by the ARC Discovery Early Career Researcher Award (DE130100868) and the Australian National Health and Medical Research Council (#1003760).

